# Application and mechanism of anticancer peptides in organoid models of intrahepatic cholangiocarcinoma

**DOI:** 10.1101/2025.02.17.638575

**Authors:** Xuekai Hu, Yue Zhang, Yanchen Li, Qiuxia Zheng, Haixia Zhao, Yun Zhang, Jingman Ni, Jia Yao

**Affiliations:** School of Pharmacy, Lanzhou University, Lanzhou, 730000, China; The First school of Clinical Medicine, Lanzhou University, Lanzhou, 730000, China; School of Basic Medical Sciences, Key Laboratory of Preclinical Study for New Drugs of Gansu Province, Lanzhou University, Lanzhou, China; The First Hospital of Lanzhou University, Lanzhou, 730000, China; Key Laboratory of Biotherapy and Regenerative Medicine, First Hospital of Lanzhou University, Lanzhou, 730000, China

**Keywords:** Intrahepatic cholangiocarcinoma, patient-derived organoid, anticancer peptide, disruption of cell membrane, apoptosis

## Abstract

**Background:** The clinical manifestations of intrahepatic cholangiocarcinoma (ICC) are non-specific, and only a small number of patients are eligible for surgical resection at the time of diagnosis, limiting treatment options. Anticancer peptides (ACPs) exhibit strong inhibitory activity against tumour cell lines, exhibit minimal side effects, are easily modified and optimised, and have low production costs. These attributes make ACPs a prospect for clinical applications. Simultaneously, the development of patient-derived three-dimensional organoids as a novel disease model has enabled the replication of the structure and heterogeneity of solid tumours. These organoids provide valuable tools for understanding disease mechanisms, conducting drug sensitivity tests, and developing targeted therapies.

However, the effect of ACPs at the organoid level in ICC remains unknown. Therefore, this study aims to explore the potential of ACPs in treating ICC using patient-derived organoids as a model system to evaluate their efficacy and optimize therapeutic strategies.

**Methods:** We designed and synthesised a novel ACP sequence and applied it to a patient-derived organoids (PDOs) model. Organoids were evaluated using histochemistry analysis and whole-exome sequencing. ACPs were compared with representative chemotherapeutic drugs, including sorafenib and gemcitabine, to assess treatment differences. Additionally, scanning electron microscopy and flow cytometry were performed for further mechanistic analyses.

**Results:** Organoid models exhibited histological characteristics similar to those of maternal tumours, especially showing strong positivity for EpCAM and KRT19 expression. Furthermore, whole-exome sequencing of the primary tumour and derived organoids revealed that PDOs closely recapitulated the genomic similarities of the primary tumour across multiple levels. Drug response analyses revealed that ACPs exhibited highly efficient anti-tumour effects, with specific differences observed between patients. ACPs affected the growth of tumour cells and exerted anticancer effects through direct membrane disruption and indirect induction of apoptosis.

**Conclusion:** In this study, organoids derived from patients with ICC were established. This *in vitro* model provides valuable tools for evaluating the therapeutic response of ACPs and offers novel insight for the study of ICC.

## BACKGROUND

Intrahepatic cholangiocarcinoma (ICC) is a rare hepatobiliary malignancy accounting for approximately 20–30% of primary liver cancers. It occurs in the liver with cholangiocyte differentiation and has limited treatment options. Moreover, it is associated with a poor prognosis. The incidence of ICC is associated with several known risk factors, including pathological conditions of the biliary system (such as cholangitis, cholestasis, and bile duct cell injury), chronic liver disease (caused by viral or non-viral infection), and metabolic abnormalities. The clinical manifestations of ICC include dull pain in the right upper abdomen, weight loss, and discomfort, with little history of jaundice. Imaging features of ICC are characterised by low-density lesions with biliary dilatation. Contrast-enhanced magnetic resonance imaging shows hypointense lesions on T1-weighted images and hyperintense lesions on T2-weighted images. Additionally, the levels of tumour markers such as glycoprotein 199 (CA199), carcinoembryonic antigen, and alpha-fetoprotein are abnormally elevated. CA199, in particular, is associated with a poor prognosis in cholangiocarcinoma [1]. Currently, surgical resection is the only treatment approach to cure ICC; however, only about 20% of patients meet the requirements for resection. Surgical resection is not recommended for patients with vascular invasion or lymph node metastasis. For patients with unresectable tumours, chemotherapy and immunotherapy are used to slow disease progression. The median survival time for patients treated with gemcitabine and cisplatin combined chemotherapy regimen is only 11.7 months [2]. Currently, no specific treatment for ICC exists, warranting the urgent development of novel drugs and innovative approaches for its management. Organoids are complex multicellular structures that, after three-dimensional (3D) conditioned culture, develop tissue-like architecture and characteristic functions. They can replicate cell-cell and cell-environment interactions. In contrast, traditional two-dimensional (2D) cell monolayer cultures lack cytoskeletal changes, enzyme activity, and cell polarity. Additionally, cells in 2D cultures tend to lose their differentiation phenotype during long-term cultivation. The construction of animal models is relatively time-consuming and labour-intensive. It retains the characteristics of animals in the microbiome, metabolome, genetics, and physiology but has certain limitations [3]. Organoid models address these shortcomings. Organoids, derived from stem cells or body tissues, provide a research model closer to human physiological and pathological conditions and simulate diseases at the tissue level. Patient-derived organoids (PDOs), typically obtained from tumour specimens through needle biopsy or surgery, can retain the histomorphological and genomic characteristics of primary tumours. These organoids have been widely used in drug screening, basic research, and translational medicine [4].

Therapeutic peptides offer several advantages, including high specificity, low toxicity, low immunogenicity, and rapid clearance, making them a promising alternative to drug-resistant chemotherapeutic agents. Anticancer peptides (ACPs) are derived from natural or modified peptides or are identified through phage display peptide library technology [5]. Appropriate modification is feasible and effective for overcoming practical obstacles such as poor stability and low permeability in the development of ACPs. Common modification methods employed include replacing neutral or anionic amino acids with cationic amino acids (such as lysine and arginine), adding chemical groups, and coupling ACPs with chemotherapeutic drugs or macromolecular polymers to enhance their targeting of cancer cells. Newly modified ACPs can potentially be transformed into drugs and vaccines. Currently, several peptide-based drugs are undergoing clinical trials to reduce disease progression and mortality. We conducted a search for ACP-related studies on the clinicalTrials.gov website and found that the included cancer types ranged from common tumours (such as melanoma, pancreatic cancer, and lung cancer) to unclassified cancers. Over 98% of these studies are interventional, while the others are observational [6]. The direct anticancer mechanism of ACPs involves forming pores in the plasma membrane, resulting in irreparable necrosis and cell lysis. The indirect mechanism of action involves dissolving internal membranes, such as mitochondrial membranes, which further triggers apoptosis. In previous studies, our research group designed and synthesised multiple ACPs based on the repetitive sequence KLLK. These ACPs demonstrate broad-spectrum anti-tumour activity *in vitro* against various cancer cell lines, including cervical cancer (HeLa), breast cancer (MCF-7), liver cancer (HepRG), and non-small cell lung cancer cells (A549). These peptides effectively inhibit the growth of tumour cells through multiple mechanisms such as cell membrane lysis, induction of cell cycle arrest, and apoptosis. However, there is no research on the application of ACPs in ICC organoids. Therefore, this study aims to employ several ACPs synthesised in the previous study to target cholangiocarcinoma cell lines and PDO models to identify alternative drugs for treating ICC [7].

## MATERIALS AND METHODS

### Peptide synthesis and purification

The peptide sequence was designed by Ms. Xiao-Yan Wu from the School of Pharmacy at Lanzhou University. It was synthesised using the standard Fmoc solid-phase synthesis method, then purified and analysed using reverse high-performance liquid chromatography (RP-HPLC; Waters, MA, USA). The molecular weight of the peptide chain was determined using electrospray ionisation mass spectrometry (ESI-MS; MaXis 4G, Bruker, USA). All samples had a purity > 95% and were stored at -20℃ in the form of lyophilised powder.

### Cell culture

The human cholangiocarcinoma cell line (HCCC-9810) and normal human intrahepatic bile duct epithelial cells (HIBEpiCs) were obtained from the Laboratory of Regenerative Medicine at the First Hospital of Lanzhou University. The cells were cultured in RPMI-1640 medium (VivaCell) supplemented with 10% foetal bovine serum (FBS; Animal Blood Ware), 100 U/mL penicillin, and 100 μg/mL streptomycin (Service Bio). The cultures were maintained in a humidified constant temperature incubator at 37 ℃ with 5% CO_2_.

### Cytotoxicity assays

The CCK-8 assay (Wilber) was used to evaluate the cytotoxicity of several peptides *in vitro*. HCCC-9810 cells were seeded into 96-well plates (NEST) at a density of 1×10^4^ cells/well and incubated for 24 h. The cells were treated with a peptide-containing medium at various concentrations for 1 h. Further, 10 μL of CCK-8 solution was added to each well, followed by incubation for an additional 4 h. A microplate reader (BioTek, Synergy H1, USA) was used to measure the absorbance at 450 nm, and the results were expressed as half-maximal inhibitory concentration (IC_50_). HIBEpiC cells were used to assess the toxicity of polypeptides on normal cells.

### Membrane potential measurement

The zeta potentials of the HCCC-9810 and HIBEpiC cells were measured using a NanoZetasizer system (Malvern, Worcestershire, UK). The peptide solution at concentrations of 0, 2.5, 5, 10, and 20 μM was incubated with a 1×10^5^ cells/mL cell suspension for 1 h. The mixture was then transferred to a folded capillary sample cell (DST1070) to measure the zeta potential value.

### Wound healing assays

HCCC-9810 cells were seeded in 6-well plates (NEST) and cultured until they reached confluence. A 200 μL pipette tip was used to create a scratch on the cell monolayer. The cells were washed with PBS (Servicebio) to remove detached cells and debris before being treated with a medium containing 1 μM polypeptide, gemcitabine (Solarbio) or sorafenib (Solarbio). Images were captured at 0 h and 24 h post-treatment using an inverted fluorescence microscope (Olympus, U-HGLGPS, Japan). The wound healing percentage was calculated based on changes in the wound area.

### Transwell assays

In Transwell-24 wells (Corning) with a pore size of 8 um, HCCC-9810 cells (1×10⁵ cells/well) were suspended in 200 μL of serum-free medium containing 1 μM polypeptide, gemcitabine or sorafenib and added to the upper chamber. The lower chamber was filled with 600 μL of RPMI-1640 medium supplemented with 10% FBS. After 24 h of incubation, the cells were washed with PBS and fixed with 4% paraformaldehyde (Solarbio) for 30 min. The cells were then washed again with PBS, stained with 0.1% crystal violet (Solarbio) for 30 min, and washed again with PBS. After removing the excess stain with a cotton swab, the sample was visualised and photographed using an inverted fluorescence microscope. The relative gray levels of each group were analysed using ImageJ software.

### Patient-derived organoid culture

This study was approved by the Ethics Committee of the First Hospital of Lanzhou University and adhered to relevant ethical guidelines (LDYYLL-2024-788). Human ICC tumour tissues were obtained from hepatectomy procedures performed in the Department of General Surgery at the First Hospital of Lanzhou University. These tissue were divided into three parts: one for organoid culture, another for histological identification, and the third for genome sequencing. Pathological examination of the intraoperative specimens showed all cases to be ICC. The tumour tissue was placed in a precooled tissue preservation solution and cut into 1 mm^3^ piece using a sterile scalpel in the dish. A small amount of digestive solution was added, and the mixture was subjected to shaking digestion at 37℃. Digestion was terminated when a sufficient number of cell clusters or single cells were observed under the microscope. The suspension was then filtered through a 100 μm mesh (Corning). The filtrate was collected and centrifuged at 300 × g for 5 min, after which the resulting supernatant was removed. The cell pellet was mixed with Matrigel (Corning) at a volume of 1:1 ratio and seeded into a 24-well plate (NEST) of 50 μL / well. After the Matrigel solidified into a gel, 500 μL of human cholangiocarcinoma organoid culture medium (Absin) was added to each well. The medium was replaced every 2–3 days. Based on the growth rate of the organoids, they were passaged at a 1:2 or 1:3 volume ratio.

### Histological evaluation of tumours and organoids

Paraffin-embedded tumour tissues and PDOs (5 μm thick) were used for histological characterisation. The prepared slides were dewaxed and rehydrated before staining. Haematoxylin and eosin (H&E) staining was performed using the H&E Kit (Solarbio), followed by microscopic examination and image acquisition. Immunohistochemistry and immunofluorescence were conducted by incubating the slides overnight with anti-KRT19 (1:200, Proteintech) and anti-EpCAM (1:500, Proteintech) antibodies. All procedures were performed according to the instructions of the manufacturer.

### Live/dead staining of organoids

The reaction buffer and staining working solution were prepared according to the instructions of the manufacturer. The organoid medium was aspirated, and the cells were gently washed with PBS by slowly adding the solution. Further, 500 μL of the staining working solution (Elabscience) was added to each well of the 24-well plate and incubated at 37℃ for 1 h. Live and dead cells were visualised using an inverted fluorescence microscope with FITC and TRITC channels, respectively. The captured images were merged and analysed using ImageJ software.

### Drug sensitivity experiment of organoids

Using a drug screening method, the response of organoids from various donors to multiple therapeutic agents was evaluated. Organoids in optimal growth conditions were embedded in 96-well plates, with 10 μL Matrigel organoid mixture per well, and cultured in a 3D environment. Bright-field images of representative wells were captured on the day treatment began. Organoids were treated with different concentrations of the compound-containing medium for 2 days. At the end of the treatment, bright-field images of the organoids from the same location were captured. The luminescence value was measured following the protocol outlined in the CellTiter-Glo kit (Promega). Wells containing only the medium and organoids served as negative controls, and the results were reported as IC_50_.

### Whole exome sequencing and data analysis

The data comprised three pairs of tumour tissues and their corresponding derived organoids. DNA was extracted from the samples and assessed for quality. The qualified DNA fragments ranged from 150–350 bp in length. After end repair and the addition of a poly-A tail, a genomic library was constructed by ligating double-ended linkers. The genomic library was hybridised in the liquid phase with biotin-labelled exome capture probes to isolate exon sequences. The target library was obtained through PCR amplification. Following quality inspection, the library was sequenced using the Illumina NovaSeq 6000 platform with PE150 sequencing.

### Scanning electron microscopy assay

Organoids in the experimental group were incubated with peptide medium at a concentration of 1× IC_50_ and incubated at 37℃ for 2 days, while those in the control group were cultured under standard conditions. Following treatment, the organoids were extracted from the Matrigel, and the resulting pellet was collected, suspended, and fixed in 3% glutaraldehyde (Lilai). After washing with ultrapure water, cells were fixed with 1% osmic acid (EMCN) for 1 h before being washed again. The samples were then dehydrated step by step with graded alcohol concentrations, dried using a Quorum (K850, UK), and the coverslips were mounted onto the sample stage using conductive glue. The mounted samples underwent gold sputter coating for conductivity enhancement (JEOL, Smart Coater, Japan). Images of the samples were acquired using a scanning electron microscope (SEM, JEOL, JSM-IT700HR, Japan).

### Nucleic acid and protein leakage assays

The leakage of cellular contents serves as an indicator of cell membrane integrity. Organoids were treated with KL-8, 14E, and 14Aad at a concentration of 1× IC_50_ for 2 days. Following treatment, the organoid culture medium was collected, and the levels of nucleic acid DNA and proteins in the secretion supernatant were measured using an ultramicro spectrophotometre (ThermoFisher Scientific, Nanadrop 2000, USA). The resulting supernatant from the organoid culture medium of the untreated group served as the negative control. Each experiment was conducted in triplicate to ensure reproducibility.

### Flow cytometry apoptosis assays

Organoids in the experimental group were incubated with a peptide-containing medium at 1× IC_50_ concentration at 37℃ for 2 days, while those in the control group were cultured under standard conditions. After removing the Matrigel from the organoids and collecting the precipitates, the organoids were digested into single cells using trypsin (Servicebio). The cells were then washed with PBS and centrifuged. Following this, the Annexin V-FITC/PI Kit (Elabscience) protocol was used to stain the cells, and they were incubated in the dark for 20 min. The level of apoptosis was measured using flow cytometry (Agilent, NovoCyte Advanteon Dx, USA).

### Statistical analysis

Statistical analysis was performed using GraphPad Prism version 10.1.2. The significance of the results was evaluated using a one-way analysis of variance (ANOVA) and two-tailed Student’s t-test. t A P -value < 0.05 was considered significant.

## RESULTS

### Peptide sequence optimization process

In our previous study, the repetitive sequence KLLK exhibited a certain degree of anti-tumour properties, especially KL-8, which shows excellent broad-spectrum anti-tumour activity. The KL-8 peptide (LLKKLLKKLLKKLL-NH_2_) was synthesised by deleting two lysine (K) residues from the N-and C-terminal ends of the KLLK sequence, resulting in a net positive charge of +7. The 14E series was developed using the amino acid scanning method to introduce glutamic acid (E) at different positions within the KL-8 sequence, with the naming scheme indicating the specific substitution site of E in the peptide chain. Substituting K with E resulted in a peptide sequence exhibiting strong haemolytic toxicity. However, replacing proline (L) with E significantly reduced haemolytic toxicity. Therefore, we continuously replaced L residues in the peptide sequence with E, resulting in a net charge of +6. Finally, the peptide chain was further modified by incorporating Aad, an analogue of E, maintaining a net charge of +6 [7]**[Figure 1a]**[8].

**Figure 1.**
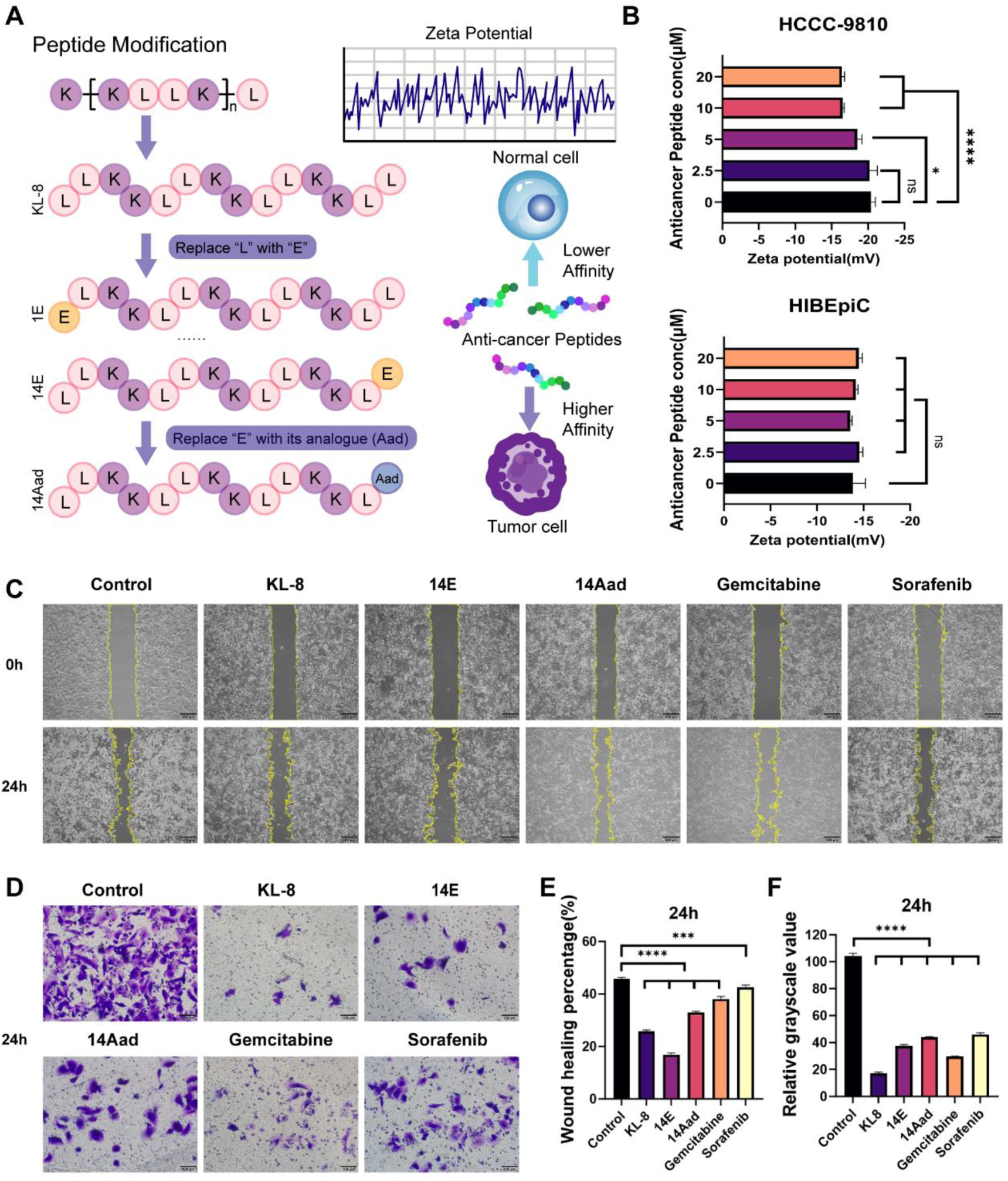
Novel anticancer peptides inhibit the proliferation of cholangiocarcinoma cells. (A) Schematic diagram illustrating peptide design. (B) Zeta potential value of cell membrane after anticancer peptides treatment. (C, E) Representative images of the scratch test showing hccc-9810 cells treated with drugs for 24 h. Scale bar, 500 μM. (D, F) Representative images of Transwell migration assay showing HCCC-9810 cells treated with drugs for 24 h. Scale bar, 200 μM. * Indicates a P value < 0.05, * * * indicates a P value < 0.001, * * * * indicates a P value < 0.0001.

### Novel ACPs inhibit the proliferation of cholangiocarcinoma cells and exhibit anticancer activity

HCCC-9810 and HIBEpiCs were selected to evaluate the anti-tumour effects of various ACPs at the cellular level. The IC_50_ values, determined through the CCK-8 assay, were used as indicators of anticancer activity. **Table 1** presents a summary of these values. Treatment with ACPs for 24 h significantly reduced the viability of HCCC-9810 cells, with peptides such as KL-8, 14E and 14Aad exhibiting IC_50_ values of ≤ 6 μM. In contrast, ACPs demonstrated lower toxicity toward HIBEpiC cells.

**Table 1.** IC_50_ values of the peptides against bile duct tumor cell and normal cell.

| Peptide | KL-8 | 1E | 2E | 5E | 6E | 9E | 10E | 13E | 14E | 14Aad |
| --- | --- | --- | --- | --- | --- | --- | --- | --- | --- | --- |
| HIBEpIC<br>IC<sub>50</sub>(μM) | 2.827 | 3.017 | 9.288 | 6.144 | 2928.00 | 69.510 | 16.230 | 18.770 | 7.583 | 6.214 |
| HCCC-9810<br>IC<sub>50</sub>(μM) | 1.686 | 8.871 | 7.025 | 9.410 | 326.00 | 24.220 | 8.356 | 10.240 | 4.506 | 5.083 |

Studies show differences in surface electrodynamic properties of normal and tumour cells after polypeptide treatment [9]. To investigate these interactions, the surface charge of the cell membrane in HCCC-9810 and HIBEpiC cells was analysed. After exposure to varying concentrations of ACPs (2.5, 5, 10, and 20 μM) for 1 h, the zeta potential of the cells was measured. For tumour cells, the zeta potential increased from -20.40 mV–16.41 mV in a concentration-dependent manner. However, no statistically significant changes in zeta potential were observed in normal cells. These findings suggest that ACPs can selectively bind to tumour cell membranes, thereby increasing the surface charge of these membranes **[Figure 1b]**.

Based on the above results, we selected three peptide sequences —KL-8, 14E, and 14Aad—for further studies [10]. Gemcitabine, a chemotherapeutic agent with proven efficacy and safety in cholangiocarcinoma treatment, and sorafenib, a first-line drug for primary liver cancer [11]. The anticancer effects of these two established chemotherapeutic drugs were compared with those of the selected peptides. The inhibitory effects of drug treatments on HCCC-9810 cells were evaluated using scratch and Transwell migration assays. All drug treatment groups demonstrated a reduction in the rate of cell migration toward the centre of the scratch after 24 h. A significant difference was observed in the wound healing rate in the experimental groups compared to that of the control group. The ACP treatment group exhibited superior inhibition of cell migration compared to that of the gemcitabine and sorafenib groups **[Figure 1c, e]**

. The Transwell migration assay revealed a significant inhibition of the migration of cholangiocarcinoma cells in the experimental group compared to that of the control group. These findings indicate that the newly designed ACPs exhibit potent anti-cholangiocarcinoma activity and effectively suppress the migration of cholangiocarcinoma cells.

### Establishment and long-term culture of organoids from ICC

Following the methodology established by Broutier et al., we obtained tumour specimens from partial hepatectomies of three patients with ICC. These specimens were successfully cultured *in vitro* to develop PDOs [12]. The primary tumour exhibited a spectrum of differentiation, ranging from low to moderate to high. **Table 2** summarises the clinical information of the patients. All three patients had a history of HBV infection, and the tumour size was > 50 cm^3^. The PDOs derived from these patients were cultured and expanded in *vitro* for over 6 months, with a passaging frequency of 1:2–1:3 every 7–14 days. The PDOs exhibited a uniform, hollow, cyst-like morphology. With the increase of culture duration, the volume and number of PDOs increased, with the larger diameters > 500 μm [13] **[Figure 2b]**.

**Figure 2.**
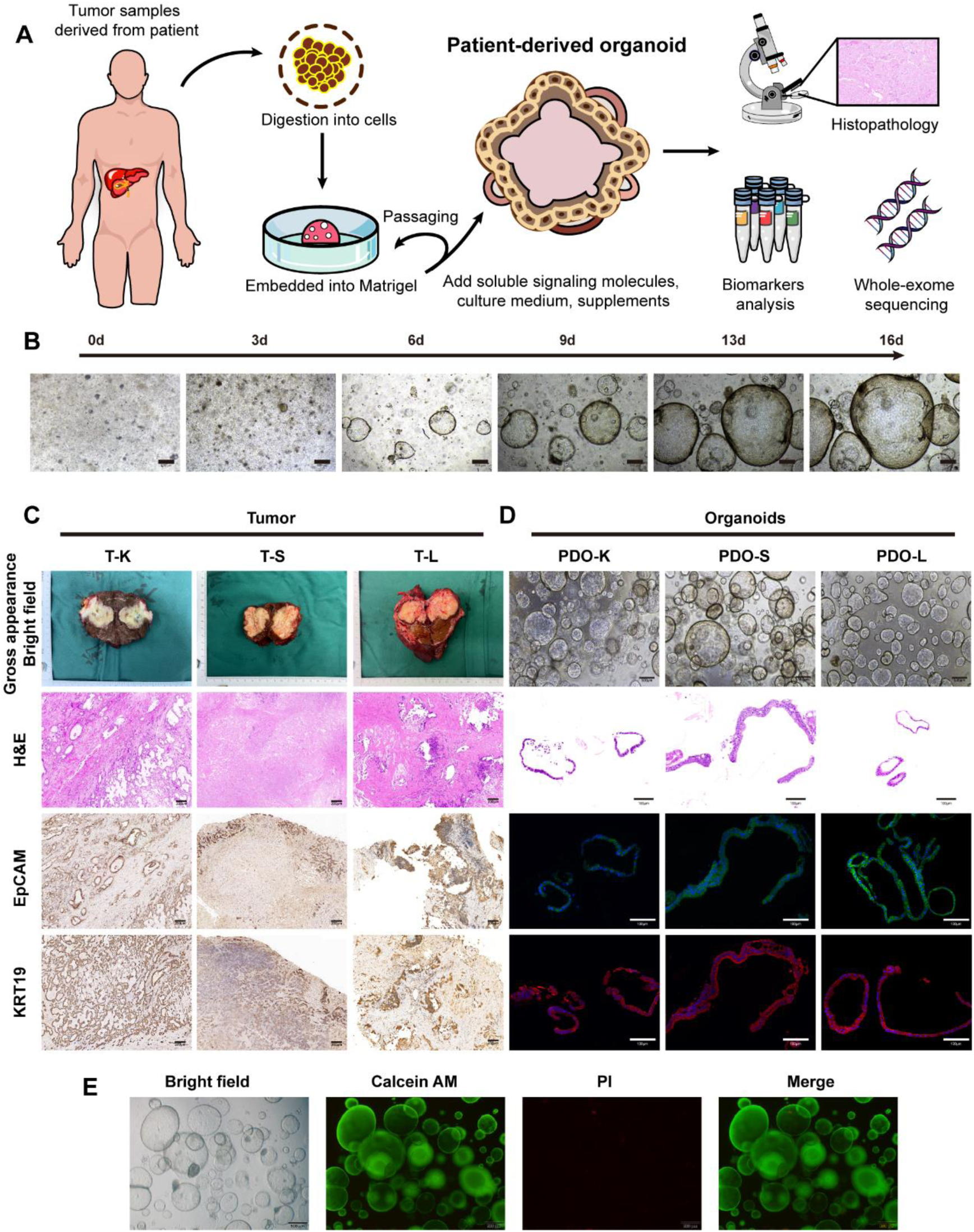
Establishment and characterization of ICC organoids. (A) Flow chart of organoid in vitro culture. (B) Morphological changes of organoids over time within 14 days. Scale bar, 500 μM. (C) Representative gross drawings, H&E staining, and immunohistochemical detection of primary tumours, including ICC universal markers (EpCAM and KRT19). Scale bar, 200 μM. (D) Bright-field diagram, H&E staining, and immunofluorescence of organoids. EpCAM (green), KRT19 (red), and nuclei were stained with DAPI (blue). Scale bar, 100 μM. (E) ICC organoids live cell (green), dead cell (red), live dead cell staining merge. Scale bar, 500 μM.

**Table 2.** Patient Data Table.

| Patient ID | Gender | Age | AFP(IU/mL) | CA199(U/mL) | CEA(ng/mL) | Virus | Tumor volume | Differentiation |
| --- | --- | --- | --- | --- | --- | --- | --- | --- |
| PDOs-K | Female | 61 | 0.87 | >1000 | 6.32 | HBV | 4x4x3.5cm | high-medium |
| PDOs-S | Male | 65 | 421.00 | 29.50 | 10.20 | HBV | 4.5x3.2x4.5cm | medium |
| PDOs-L | Female | 60 | 2.83 | 9.95 | 0.82 | HBV | 8.5x4.5x4cm | medium-low |

H&E staining, immunohistochemistry, and immunofluorescence were used to characterise the tumour tissues and derived organoids. H&E staining revealed that the organoids exhibited irregular cyst-like and adenoid shapes, similar to the original tumour tissue structure. Epithelial cell adhesion molecule (EpCAM), a cell surface glycoprotein extensively expressed in several epithelial cells, plays a critical role in cell adhesion, signalling, and differentiation. Keratin 19 (KRT19) is an important intermediate filament protein primarily expressed in epithelial cells. Overexpression of EpCAM and KRT19 is associated with ICC and is considered a potential biomarker for this disease. In patients with ICC, high expression levels of EpCAM and KRT19 correlate with increased tumour aggressiveness and poor prognosis [14, 15]. These proteins serve as potential diagnostic markers and therapeutic targets, providing valuable insight into the biological characteristics of ICC. EpCAM and KRT19 were highly expressed in all tumour tissues and derived organoids. This indicates that the expression pattern of organoids aligns with those observed in patient tissues [16] **[Figure 2c and d]**.

Organoids are three-dimensional structures composed of multiple cell types. Using fluorescence staining to differentiate between live and dead PDO cells, the proliferation and apoptosis states of the organoids can be visually observed [17]. This staining process also revealed that the organoids exhibited hollow, cystic structures, with most cells in a viable state. These findings indicate that the cultured PDOs maintain high cell viability **[Figure 2e]**.

In general, PDOs retain the tissue structure of the original specimen and exhibit high viability even after long-term culture and expansion.

### Genetic characteristics of PDOs

We performed whole-exome sequencing on the three organoids and their corresponding parental tumours to assess whether the organoids retained the genome of the parental tumour [18]. The results showed that tumour tissues and their derived organoids from the same patient shared the majority of variant sites **[Figure 3a]**. Furthermore, all three sample groups consistently expressed AHNAK, PDE4DIP, PABPC1, and BCLAF1. Studies show that these loci affect the occurrence and progression of ICC or cholangiocarcinoma [19–22]. Besides commonly observed ICC-related mutation sites, each patient exhibited unique mutation sites specific to their disease status. For example, T_S and PDO_S express ubiquitin-specific peptidase 8 (USP8), which is associated with poor prognosis in patients with ICC. USP8 enhances ICC malignancy by stabilising O-GlcNAc transferase through its deubiquitination activity [23]. Human leukocyte antigen-A, a critical component of highly immunogenic ICC, is moderately and strongly expressed in approximately 66% of ICC cases [24]. This mutation was also detected in the T_K and PDO_K samples. Additionally, T_L and PDO_L exhibited more mutations associated with poor prognosis, including COL7A1, DNMT3A, FCGBP, and IDH1among others [25–28]. These results show that the PDOs established *in vitro* effectively retain the genetic characteristics of the parental tumours and reproduce the mutation profile unique to individual patients.

**Figure 3.**
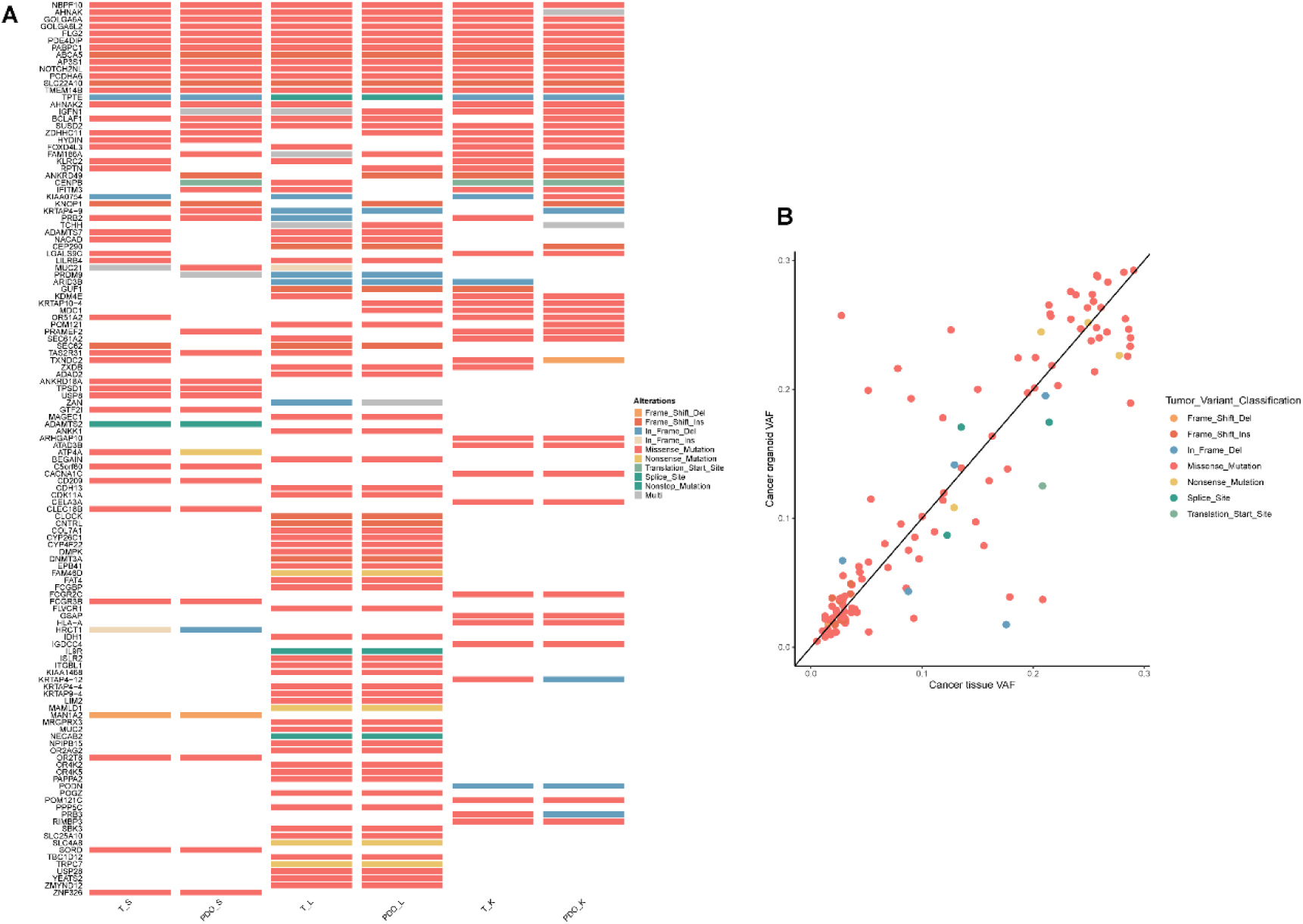
WES of patient-derived organoids and corresponding parental tumour. (A) Analysis of the SNVS of organoids and matched primary tissues shows the mutated gene type. (B) Scatter plot displaying the variant allele frequencies of shared mutations among corresponding samples.

### Anticancer peptide drug sensitivity study of established PDOs

Different dilution series of ACPs, gemcitabine, and sorafenib were added to the organoid culture medium to evaluate the specific response of patients to the drug. After the intervention, the viability of 3D cultured cells was determined by measuring ATP levels, and drug sensitivity was expressed as the IC_50_ value [29] **[Figure 4a]**. PDOs in the experimental group exhibited significant morphological changes, including shrinkage, reduced volume, and loss of their original three-dimensional morphology. The organoid changed colour from transparent to dark and cloudy, with some areas showing black necrotic regions. In the 3D culture system, the overall structure of the organoids became loose, often detaching from the matrix. Additionally, the matrigel dome structure was affected, losing its ability to maintain a stable three-dimensional structure. In contrast, the control group showed no morphological changes, continued to grow, and maintained strong viability **[Figure 4c]**.

**Figure 4.**
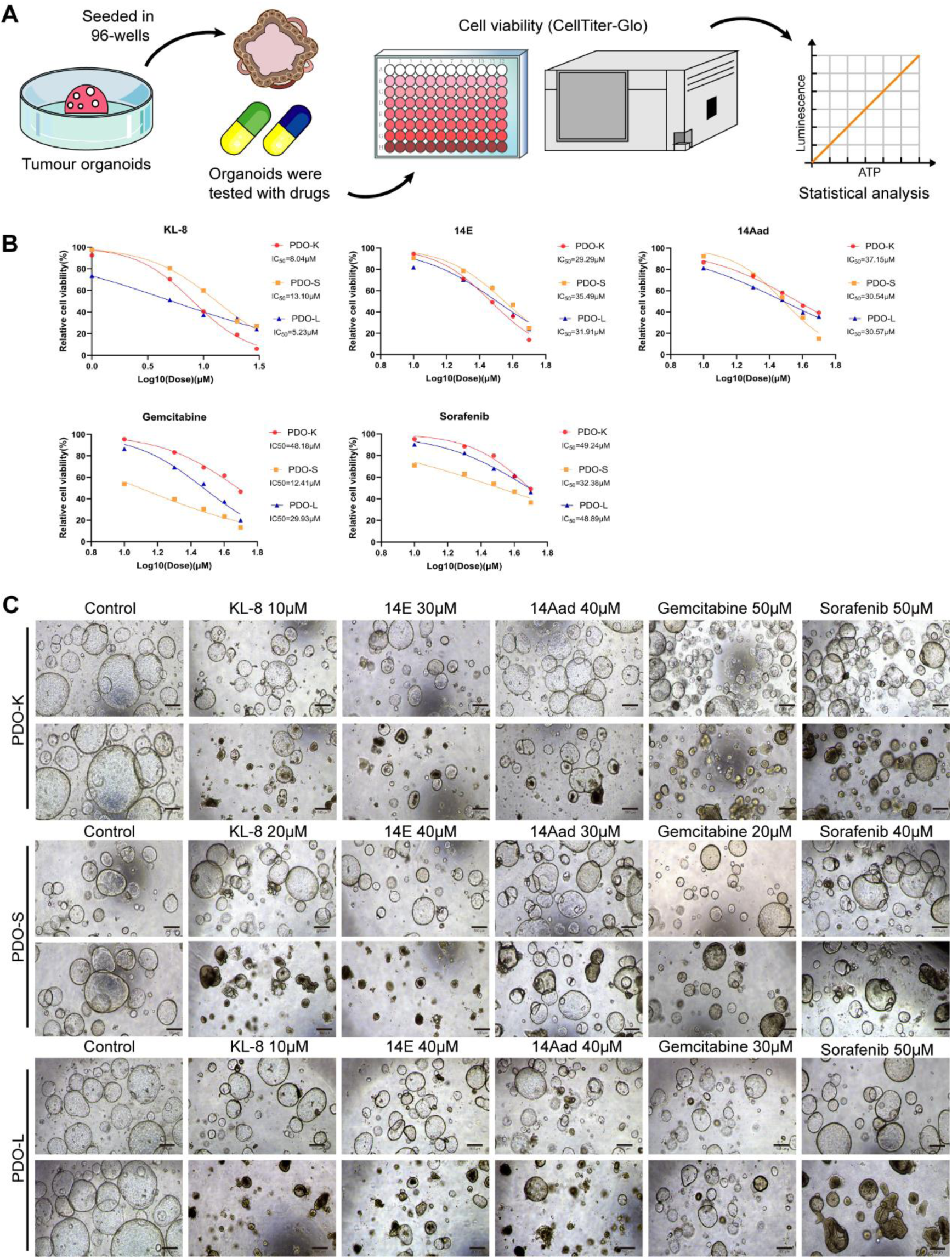
Response of patient-derived tumour organoids to drug treatment. (A) Schematic diagram of the drug sensitivity test. (B) The IC_50_ values of three organoids for different drugs were calculated. (C) Representative bright field diagram of organoids after drug action. Scale bar, 500 μM.

We observed that, compared to that of the IC_50_ values obtained for the HCCC-9810 cell line, the IC_50_ values for each group of organoids were higher. These findings indicate that the 3D organoid model more closely replicates the *in vivo* environment compared to that of the 2D cell culture, offering more accurate drug sensitivity results [30, 31]. However, individual differences in drug sensitivity were observed among the patients. KL-8 demonstrated the best therapeutic effect on PDO-K and PDO-L, while gemcitabine was most effective for PDO-S. Overall, the three PDOs exhibited the highest sensitivity to KL-8 and the lowest sensitivity to sorafenib **[Figure 4b]**. PDOs demonstrated potential as a valuable drug screening tool with varying responses to different drugs. Additionally, ACPs showed significant therapeutic effects for ICC treatment, which are comparable to or even superior to those of traditional chemotherapy drugs.

### Anti-tumour mechanism of ACPs on ICC

Studies show that ACPs can disrupt and lyse cell membranes through interactions with the cells. To further investigate this mechanism, we employed SEM imaging to observe the morphological changes in the organoids after drug intervention. PDOs in the control group showed tight junctions between cells, forming aggregated cell clumps. The overall morphology of the cell surface was smooth, continuous, and well-defined, displaying a refined microstructure. The cell surface contains microvilli, which enhance surface area and facilitate absorption and excretion [32]. Cilia are involved in cell movement, sensing the external environment, and facilitating cell-cell communication [33]. Additionally, cell protrusions contribute to the roughness of the cell surface. Owing to the action of the ACPs, the surfaces of the PDOs became rough. Obvious voids and content exudation were observed in single cells. Simultaneously, microvilli and cilia appeared to detach or break, indicating significant cell damage **[Figure 5a]**. We collected the culture medium from PDOs treated with varying concentrations of ACPs to measure the release of nucleic acid DNA and protein. The results showed that all three ACPs significantly increased the levels of released nucleic acid DNA and protein. However, no notable differences were observed between the different concentrations tested (data not shown) **[Figure 5b]**. These findings indicate that ACPs effectively disrupt cell membranes, resulting in the leakage of nucleic acid and protein from the cells. This suggests that ACPs directly induce tumour cell death through rapid membrane lysis.

**Figure 5.**
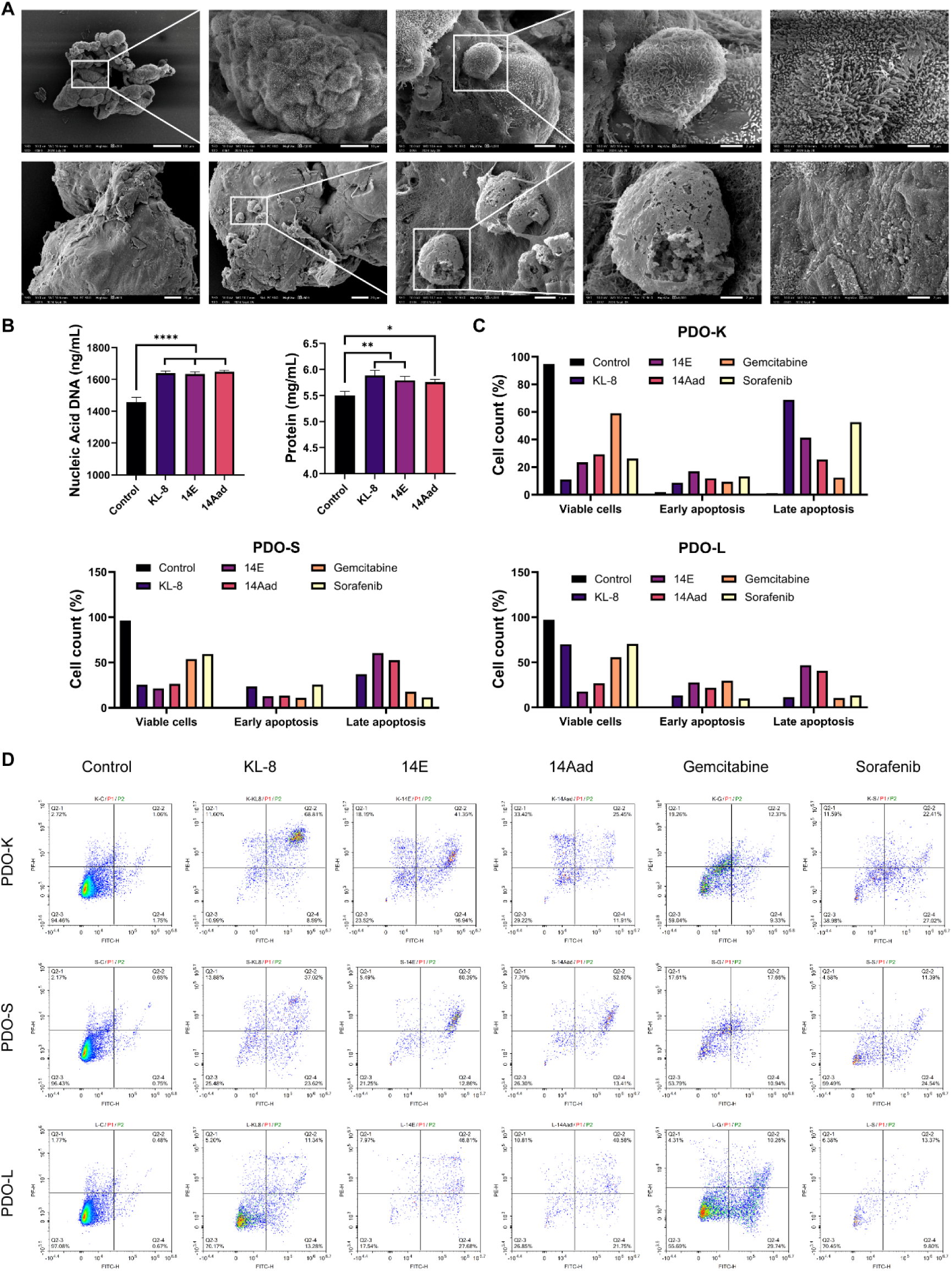
Anti-tumour mechanism of anticancer peptides. (A) Morphological changes of organoids after anticancer peptides treatment. (B) After exposure to anticancer peptides, nucleic acids and proteins in organoid culture medium leak** Indicates a P value <0.01, * * * indicates a P value < 0.001, * * * * indicates a P value < 0.0001. (C, D) organoids were treated with anticancer peptides for 24 h and digested into single cells for apoptosis assays.

Besides their direct membrane-disrupting mechanisms, ACPs can also induce apoptosis. PDOs were treated with 1× IC_50_ concentrations of the respective intervention drugs for 24 h, after which cells were digested into single-cell suspensions using trypsin and analysed for apoptosis via flow cytometry. Compared to that of the control group, the experimental group induced apoptosis to different degrees **[Figure 5c and d]**. These findings indicate that the proliferative activity of ACPs against cholangiocarcinoma cells is also associated with apoptosis.

In summary, ACPs disrupt the integrity of the cell membrane, resulting in cell death through direct interaction with the membrane. Moreover, it plays an anti-tumour role by promoting cancer cell apoptosis.

## DISCUSSION

Although significant progress has been made in the prevention, diagnosis, and treatment of ICC in recent years, no new drug types have been discovered. Studies show that therapeutic peptides have been developed for the treatment of various diseases, including cancer, microbial infections, and metabolic disorders. ACPs have emerged as a class of potential drug candidates owing to their broad-spectrum anticancer activity and high biocompatibility. Adjustments to physical and chemical properties have been implemented to optimise ACPs, including modifications to their conformation, net charge, hydrophobicity, and secondary structure [34]. The optimal peptide identified in this study showed a high α-helix structure with appropriate hydrophobicity and ideal modification sites. It demonstrated a rapid killing effect within 30 min of application. Regarding anti-tumour mechanisms, cancer and normal cells show obvious differences in appearance, genome stability, growth and reproduction, angiogenesis, and other characteristics. Cancer cells are characterised by unlimited proliferation, immune system evasion, metastasis, and invasion. The cell membrane resists adverse cellular environments and plays an important role in various anticancer mechanisms. The ACPs designed and synthesised had a higher tendency to bind to cancer cell membranes. The possible reasons for this result are as follows: the cell membrane consists of lipids and proteins, with phosphatidylserine (PS) being a key component, which is negatively charged. The PS of cancer cells is located in the outer layer of the membrane, resulting in a negatively charged surface. Additionally, other anionic components, such as O-glycosylated mucin, sialylated gangliosides, and heparan sulphate, contribute to the negative charge of the cancer cell membrane. In normal cells, amphoteric phosphatidylcholine and sphingomyelin are present in the outer layer of the plasma membrane, typically maintaining a neutral charge. However, the tumour microenvironment, characterised by high levels of reactive oxygen species and hypoxia, disrupts the balanced distribution of outer membrane components. This imbalance can lead to an increased presence of anionic molecules, resulting in a higher negative charge on the cell membrane [35, 36]. Cancer cells have numerous microvilli on their outer membrane, resulting in a larger surface area and enhancing membrane fluidity [37]. This structural difference in the cell membrane allows our ACPs to exhibit partial selectivity for tumour cells, which is a potential therapeutic method.

Organoids are promising preclinical models with significant potential for evaluating the effects of drug treatments. Tumour organoids can accurately summarise some important characteristics of solid tumours, including their internal structure, cellular heterogeneity, signalling pathways, cell-cell interactions, growth dynamics, gene expression profiles, and drug resistance mechanisms [38, 39]. In this study, we successfully established three ICC organoid models using clinical samples obtained from surgical procedures. These organoids demonstrated stable and long-term expansion capabilities. Histological and genetic characterisations confirm that the PDOs were derived from tumour tissue and maintained consistency with their parental tumour. Further, *in vitro* drug evaluation revealed that the organoids exhibited patient-specific heterogeneity, with varying drug sensitivities among patients. This 3D model allowed us to observe the effective inhibition of ICC by ACPs and explore its potential anticancer mechanism. Scanning electron microscopy revealed detailed insight into the sub-microscopic structure of the organoids. The cells on the surface of the organoids were tightly connected and formed compact clusters. The cell surfaces featured functional microvilli and cilia, which provided structural support and facilitated cellular interactions. Upon ACP treatment, the integrity of the cell membrane was compromised, leading to visible damage and leakage of intracellular contents. Additionally, cilia and microvilli detached or broke across an extensive area, affecting the movement and absorption functions of tumour cells, thereby inhibiting tumour growth and migration. ACPs eliminate cancer cells primarily by disrupting the integrity of the cell membranes. Additionally, beyond this direct mechanism of membrane destruction, ACPs also destabilise cancer cells indirectly by inducing apoptosis.

The main limitation of this study was its small sample size, which prevents the determination of the preference and correlation of ACPs in drug treatment effects on patients with ICC having different pathological types. Additionally, the limited number of organoids and lack of standardisation culture protocols restricted the ability to perform high-throughput drug screening. Transitioning to automated organoid culture procedures could significantly enhance screening efficiency [40]. Additionally, our culture system lacks a tumour microenvironment, such as immune cells, fibroblasts, and vascular system, which limits its ability to accurately predict immunotherapy outcomes. In future experiments, immune system-related cells could be incorporated into co-culture models to address this limitation [41]. Finally, our ACPs have not been clinically tested; hence, the data obtained from organoid experiments cannot yet provide patients with reliable references for clinical medication. The clinical application of ACPs in combination with PDOs will require extensive experimental validation [42].

## CONCLUSIONS

Our results provide strong evidence for the effectiveness of ACPs in ICC organoids. The study showed novel designed ACPs inhibited the progression of ICC by destroying the cell membrane and inducing apoptosis. These findings suggest that ACPs may be effective candidate compounds for the treatment of ICC in the future, offering novel ideas for therapeutic development.

## LIST OF ABBREVIATIONS

ICC: intrahepatic cholangiocarcinoma
ACPs: anticancer peptides
PDO: patient-derived organoid
3D: three-dimensional
2D: three-dimensional
HCCC-9810: human cholangiocarcinoma cell line
FBS: foetal bovine serum
H&E: Haematoxylin and eosin
EpCAM: Epithelial cell adhesion molecule
KRT19: Keratin 19
USP8: ubiquitin-specific peptidase 8
PS: phosphatidylserine

## DECLARATIONS

### Ethics approval and consent to participate

The study was conducted according to the guidelines of the Declaration of Helsinki and approved by the Ethics Committee of Lanzhou University First Hospital (2023 A-381).

### Consent for publication

Not applicable.

### Availability of data and materials

The datasets used and/or analysed during the current study are available from the corresponding author on reasonable request.

### Competing interest

The authors declare that they have no competing interests.

### Funding

This work was supported by Grant(s) from the National Natural Science Foundation of China (32160230, 82173678, 81773564), Talent Innovation and Entrepreneurship Project in Chengguan District, Lanzhou City, Gansu Province (2022RCCX0024), and 1st Hospital of Lanzhou University Scientifc Research Foundation (ldyyyn2019-03).

### Authors’ contributions

Y. Z, Y.C.L, and Q.X.Z for cell culture, X.K.H and H.X.Z was responsible for specimen collection, and they were the main contributors to writing the manuscript. X.K.H cultured and identified organoids. Y. Z, Y.C.L, and Q.X.Z and H.X.Z modifies the image and participates in data processing. X.K.H, Y.Z, Y.C.L, Q.X.Z, H.X.Z and Y.Z organized the data of all experiments. J.M.N and J.Y conceived, designed, and supervised the study. All authors read and approved the final manuscript.

## Acknowledgements

We sincerely appreciate all the participants in our work.

## Authors’ information

School of Pharmacy, Lanzhou University, Lanzhou, 730000, China

Xuekai Hu, Yun Zhang & Jingman Ni

The First school of Clinical Medicine, Lanzhou University, Lanzhou, 730000, China

Yue Zhang, Yanchen Li, Qiuxia Zheng, Haixia Zhao & Jia Yao

School of Basic Medical Sciences, Key Laboratory of Preclinical Study for New Drugs of Gansu Province, Lanzhou University, Lanzhou, China

Yun Zhang & Jingman Ni

The First Hospital of Lanzhou University, Lanzhou, 730000, China

Jia Yao

Key Laboratory of Biotherapy and Regenerative Medicine, First Hospital of Lanzhou University, Lanzhou, 730000, China

Jia Yao

## Notes

### Competing Interest Statement

The authors have declared no competing interest.

